# Gradients of structure-function tethering across neocortex

**DOI:** 10.1101/561985

**Authors:** Bertha Vázquez-Rodríguez, Laura E. Suárez, Golia Shafiei, Ross D. Markello, Casey Paquola, Patric Hagmann, Martijn P. van den Heuvel, Boris C. Bernhardt, R. Nathan Spreng, Bratislav Mišić

**Affiliations:** McConnell Brain Imaging Centre, Montréal Neurological Institute, McGill University, Montréal, Quebec, Canada; Signal Processing Laboratory (LTS5) Ecole Polytechnique Fédérale de Lausanne, Route Cantonale, 1015 Lausanne, Switzerland; Center for Neurogenomics and Cognitive Research, Vrije Universiteit Amsterdam, The Netherlands

## Abstract

The white matter architecture of brain networks imparts a distinct signature on neuronal co-activation patterns. Inter-regional projections promote synchrony among distant neuronal populations, giving rise to richly patterned functional networks. A variety of statistical, communication and biophysical models have been proposed to study the relationship between brain structure and function, but the link is not yet known. In the present report we seek to relate the structural and functional connection profiles of individual brain areas. We apply a simple multilinear model that incorporates information about spatial proximity, routing and diffusion between brain regions to predict their functional connectivity. We find that structure-function relationships vary markedly across the neocortex. Structure and function correspond closely in unimodal, primary sensory and motor regions, but diverge in transmodal cortex, corresponding to the default mode and salience networks. The divergence between structure and function systematically follows functional and cytoarchitectonic hierarchies. Altogether, the present results demonstrate that structural and functional networks do not align uniformly across the brain, but gradually uncouple in higher-order polysensory areas.

## INTRODUCTION

Intricate connection patterns among neural elements form a complex hierarchical network that promotes signaling and molecular transport [10, 58]. Neural elements have a pronounced tendency to form local cliques and tightly-coupled communities with common functional properties [29]; a small proportion of long-distance projections allows signals to be sampled and integrated from these specialized domains [5, 37, 74]. Perpetual interactions via the white matter “connectome” manifest as richly patterned neural activity and are thought to support perception, cognition and action [52].

What is the link between structure and function in brain networks? Relating the organization of physical connections to patterns of functional interactions is a key question in systems neuroscience. A number of methods have been used to address this link, including statistical models [44, 47], communication models [14, 26, 49] and biophysical models [8, 19, 31, 65]. The focus has traditionally been on using whole-brain structural connectivity to predict whole-brain functional connectivity, with the assumption that a common mechanism operates across the entire network. These methods have proven insightful and generally yield moderate fits to empirical functional connectivity patterns, from approximately 25% to 50% of the variance explained [45].

Nevertheless, structure and function may not be related in exactly the same way across the whole brain. Recent evidence points to a fundamental organizing principle for macroscale functional interactions [42]. The hierarchy spans from unimodal primary areas to polysensory transmodal areas, tracing a continuous sensory-fugal gradient that culminates in the default mode network [36, 46]. This representational gradient may reflect microstructural variations, showing significant associations with intracortical myelination [33] and laminar differentiation [56]. Altogether, this work opens the possibility that structure and function may not be related in exactly the same way across the whole brain, but potentially converge or diverge in specific areas.

Here we address the relationship between structure and function by focusing on connection profiles of individual brain regions. We first reconstruct structural and functional networks from diffusion MRI (dMRI) and resting-state functional MRI (fMRI) in a cohort of 40 healthy participants. We then apply a simple multilinear model that uses information about a region’s geometric and structural network embedding to predict its functional network embedding. The method allows us to ask how closely structure and function correspond in individual regions and the extent to which this correspondence reflects affiliation with cognitive systems, cytoarchitecture and functional hierarchies.

## RESULTS

Structural and functional networks were reconstructed as follows:

- *Structural networks*. Structural and functional connectivity were derived from *N* = 40 healthy control participants (source: Lausanne University Hospital). Structural connectivity was estimated from diffusion spectrum imaging. Adjacency matrices were reconstructed using deterministic streamline tractography. A group-consensus structural connectivity matrix was assembled using a consistency- and length-based procedure [4, 6, 48, 49].
- *Functional networks*. Functional connectivity was estimated in the same healthy individuals using resting-state functional MRI (rs-fMRI). Functional connections were defined as zero-lag Pearson correlations among regional time courses. A group-consensus functional connectivity matrix was estimated as the mean connectivity of stable pair-wise connections across individuals (see *Materials and Methods* for more information on the procedure).

Initial data exploration was performed at the highest resolution (1000 nodes), using group consensus structural and functional networks (see *Materials and Methods* for more details). Analyses were subsequently repeated at other resolutions and for individual participants, and in an independently-collected dataset.

To estimate the correspondence between local structure and function, we constructed a multilinear regression model that relates node-wise structural and functional connectional profiles (Fig. 1). For a given node *i*, the dependent variable is the resting state functional connectivity between node *i* and all other nodes in the network *j ≠ i*. The predictor variables are the geometric and structural relationships between *i* and *j*, including Euclidean distance, path length and communicability. The observations or samples are the individual *i, j* relationships. Model parameters (regression coefficients for each of the 3 predictors) are then estimated via ordinary least squares. Goodness-of-fit for each node *i*, representing the correspondence between structural and functional profiles for that node, is quantified by the adjusted 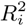 between observed and predicted functional connectivity. The use of a multilinear model to relate structure and function is conceptually similar to the method previously reported by Gonñi and colleagues [26] (see also [7, 44]), with the important exception that the present model focuses on connection profiles of individual regions rather than whole-brain connectivity.

**Figure 1.**
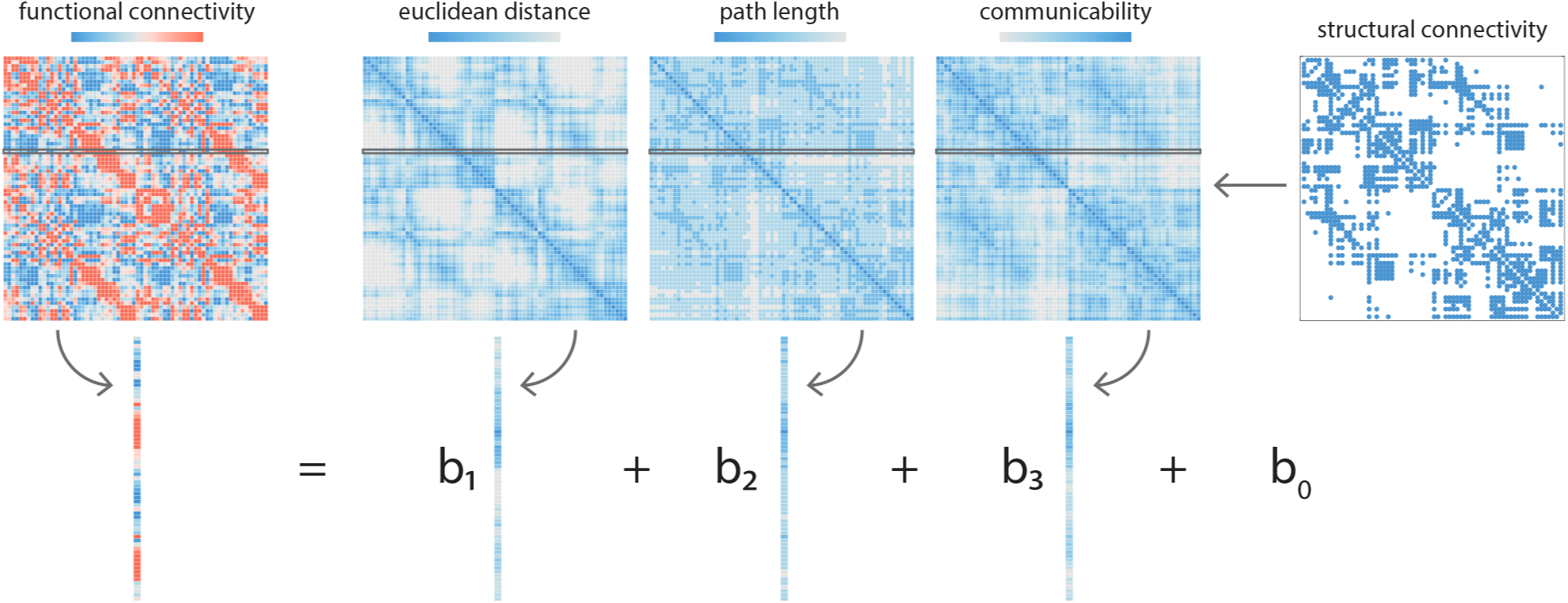
Node-wise structure-function relationships. Local, node-wise structure-function relationships are estimated by fitting a multilinear regression model for each node separately. For a given node *i*, the response or dependent variable is the functional connectivity between node *i* and node *j* ∆ *i*. The predictor or independent variables are the geometric and structural relationships between *i* and *j*, including the Euclidean distance, path length and communicability. The “observations” are individual *i, j* relationships. Model parameters (regression coefficients *b*_1_, *b*_2_ and *b*_3_) are then estimated by ordinary least squares. Goodness-of-fit for each node *i* is quantified by 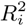 between observed and predicted functional connectivity.

### Convergent and divergent structure-function relationships across neocortex

The correspondence between structural and functional connection profiles is highly variable across neocortex. Fig. 2a shows the histogram of *R*^2^ values from each of the node-wise multilinear models. Mean *R*^2^ = 0.30 (median *R*^2^ = 0.30), roughly concordant with previous reports that used similar models to predict whole-network functional connectivity [26]. However, the values vary considerably, from *R*^2^ = 0.04 to *R*^2^ = 0.62 (inter-quartile range = 0.18), indicating that for some regions there is a strong correspondence between structural network embedding and function, while for others there is little evidence of any such correspondence.

**Figure 2.**
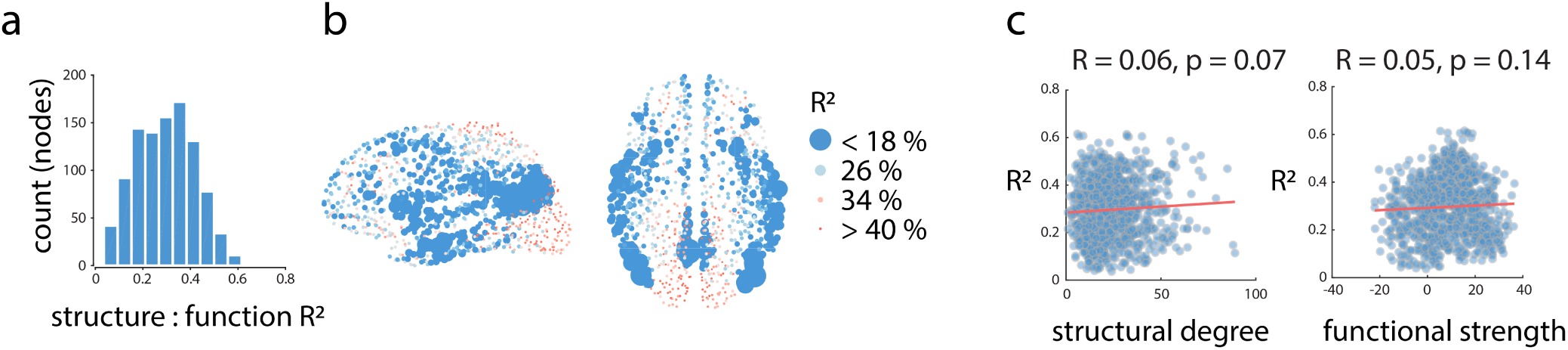
Convergent and divergent structure-function relationships across neocortex. (a) Local structure-function correspondence, estimated by node-wise *R*^2^ from the multilinear model. The histogram shows a wide distribution of *R*^2^ values across 1000 nodes at the highest resolution. (b) The spatial distribution of structure-function correspondence. Nodes are coloured and sized in inverse proportion to *R*^2^; nodes with weaker structure-function correspondence are larger. High correspondence is observed in primary sensory and motor cortices, while lower correspondence is observed in transmodal cortex. (c) Correlation between structural and functional centrality and structure-function correspondence. Scatter plots between node-wise *R*^2^ and structural and functional centrality, estimated by binary degree and weighted strength, respectively. The low correlations suggest that the correspondence between structure and function does not trivially depend on the structural or functional connectedness of a node. For the same results at other parcellation resolutions, please see Fig. S1.

We next examine the anatomical distribution of structure-function *R*^2^ values. To highlight regions that show little correspondence, node size and colour are inversely proportional to their *R*^2^ (Fig. 2b). The map shows a highly organized and hemispherically symmetric spatial arrangement. Brain regions with least structure-function correspondence include medial parietal structures (precuneus, posterior cingulate), lateral parietal and temporal cortices, insular cortex and anterior cingulate cortex. Conversely, primary sensory regions, including occipital and paracentral cortices show relatively high structure-function correspondence.

It is possible that low *R*^2^ values are observed in some areas because they have either too many or too few direct connections. To examine this possibility, we correlated regional *R*^2^ values with the structural degree and functional strength of each node (Fig. 2c). In both cases the correlations were low (structural: *R* = 0.06, *p* = 0.07; functional: *R* = 0.05, *p* = 0.14), suggesting that regional variations in structure-function correspondence were not trivially driven by structural or functional centrality. We subsequently repeated these analyses for all 5 resolutions of the Lausanne atlas. The results are shown in Fig. S1 and are consistent across resolutions. We also replicated these findings in an independently-collected dataset at resolutions 2, 3 and 4 (Human Connectome Project; Fig. S2). The spatial patterns of *R*^2^ values are visually similiar (Fig. S2a) and significantly correlated (R = 0.77, 0.72 and 0.67; Fig. S2b).

### Structure-function relationships follow functional and cytoarchitectonic hierarchies

The spatial distribution of *R*^2^ values suggests that structure-function correspondence may be circumscribed by functional systems or cytoarchitectonic attributes. To address this question, we applied two partitions: (1) resting state networks described by Yeo and colleagues [84], (2) cytoarchitectonic classes described by von Economo and Koskinas [66, 77, 80]. The former groups brain regions according to how similar their time courses are and the latter groups regions according to how similar they are in terms of cell morphology.

We first calculated the mean *R*^2^ for each network or class. To assess the extent to which these means are determined by the partition, and not trivial differences in size, coverage or symmetry, we used a label-permuting null model [49]. Network or class labels were randomly re-assigned and mean *R*^2^ values were re-computed (1,000 repetitions). The network- or class-specific mean *R*^2^ was then expressed as a z-score relative to this null distribution.

There is a gradual divergence between structure and function moving from unimodal to transmodal cortex. Fig. 3 shows the z-scored *R*^2^ for each resting state network (red) and cytoarchitectonic class (blue). Positive values indicate that the structure-function relationship is stronger than expected by chance, while negative values indicate that the structure-function relationship is weaker than expected by chance. Consistent with the intuition developed in the previous section, statistically significant divergence between structure and function is observed in polysensory or transmodal cortex, namely the default mode and ventral attention resting state networks, and the insular and association classes. The reverse is true for primary unimodal cortex, where there is a significant convergence between structure and function.

**Figure 3.**
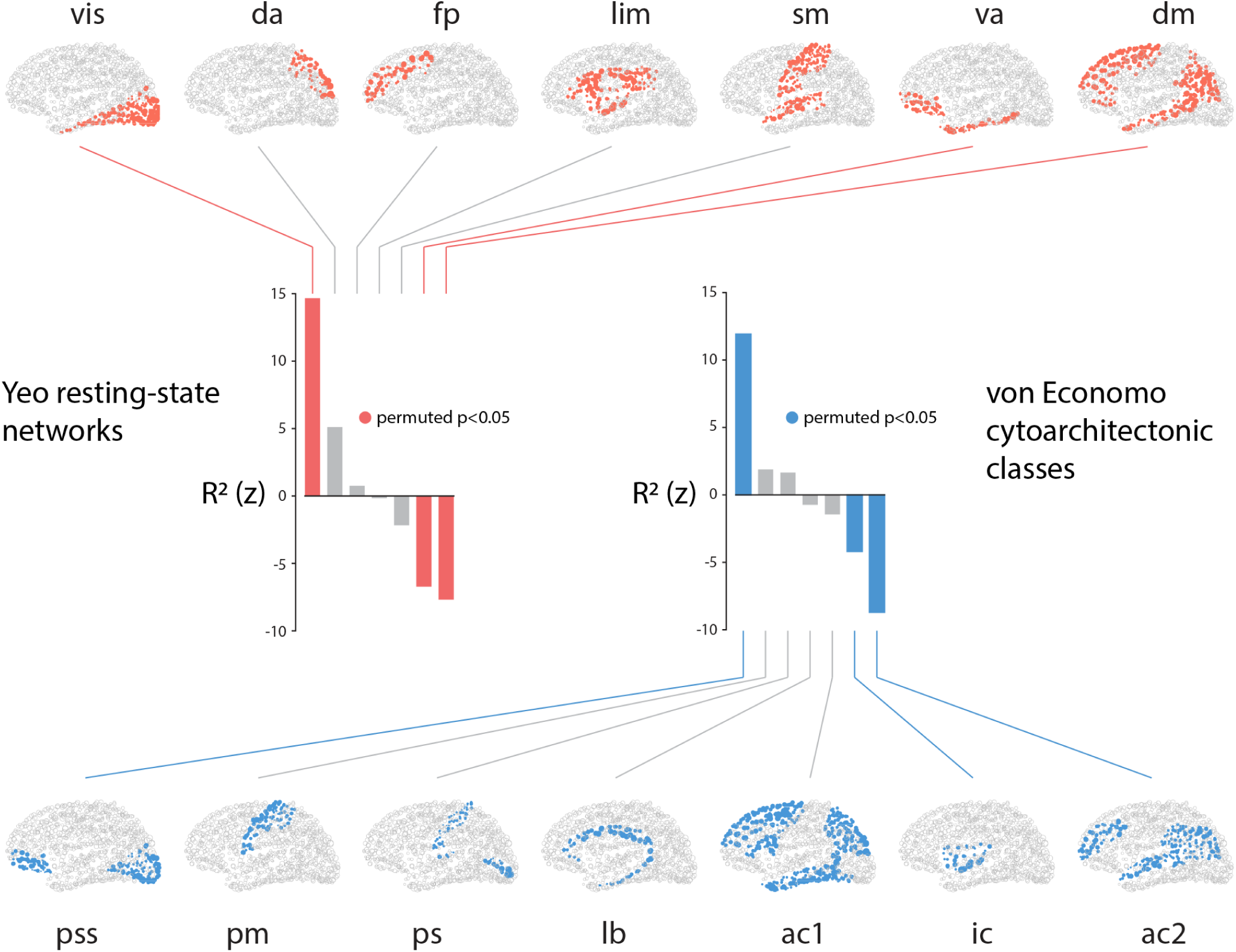
Structure-function tethering across cognitive systems and cytoarchitectonic classes. Node-wise *R*^2^ values are averaged according to their membership in resting-state networks or cytoarchitectonic classes. To determine whether the mean value for each network or class is statistically significant, a null distribution is constructed by randomly permuting the network or class label (10,000 repetitions). The network- or class-specific mean *R*^2^ is then expressed as a z-score relative to this null distribution. Statistically significant networks/classes are shown in colour; non-significant networks/classes are shown in grey. Yeo networks: vis = visual, da = dorsal attention, fp = frontoparietal, lim = limbic, sm = somatomotor, va = ventral attention, dm = default mode. von Economo classes: ac1 = association cortex, ac2 = association cortex, pm = primary motor cortex, ps = primary sensory cortex, pss = primary/secondary sensory, ic = insular cortex, lb = limbic regions.

### Structure and function systematically diverge along a macroscale functional gradient

Recent studies suggest a universal organizational principle whereby brain regions are situated along a continuous gradient or hierarchy, ranging from primary sensory and motor regions to transmodal regions [34, 42]. It is therefore possible that the patterns of structure-function convergence and divergence recapitulate this hierarchy.

We first derived a macroscale functional gradient for the present data set. The correlation-based functional network was converted to a transition probability matrix and subjected to singular value decomposition, a method known as diffusion map embedding [13] (see *Materials and Methods* for more details). The first eigenvector of the matrix, which we refer to as a “gradient”, spans primary unimodal cortex on one end and transmodal cortex on the other (Fig. 4a). Critically, the map bears a strong resemblance to the vertex-wise map originally reported by Margulies and colleagues [42].

**Figure 4.**
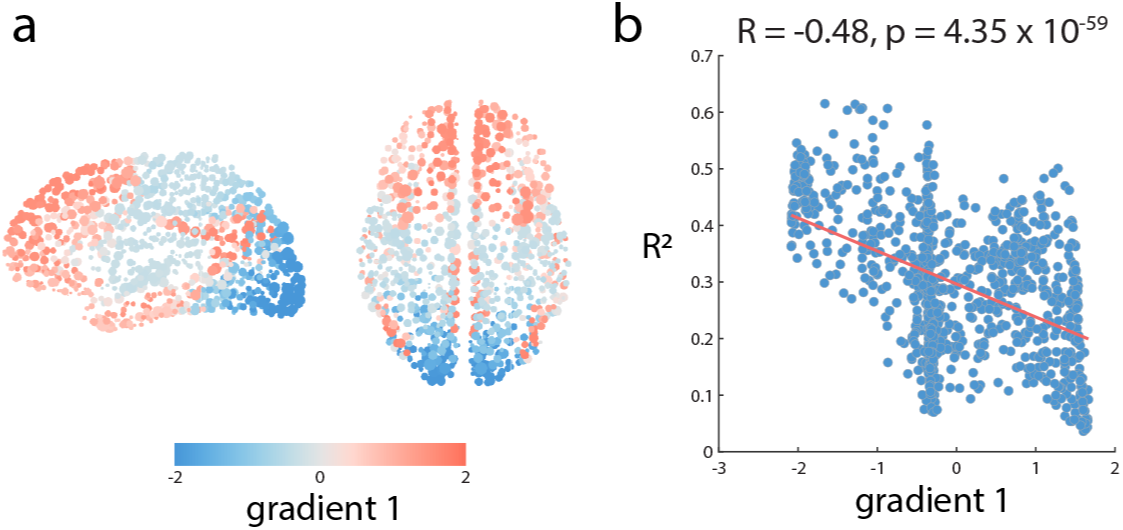
Structure-function divergence across large-scale functional network gradients. Large-scale functional network gradients were identified by applying diffusion map embedding to the normalized graph Laplacian of the correlation matrix. (a) The first gradient runs from primary, unimodal cortex to transmodal cortex and resembles the vertex-wise map originally reported by Margulies and colleagues [42]. (b) Node-wise structure-function *R*^2^ values are anti-correlated with positions along this gradient, suggesting that structure and function closely correspond in unimodal cortex but diverge in transmodal cortex.

We then assess the relationship between structure-function *R*^2^ for a given region, and its position along the macroscale functional gradient. Fig. 4b shows that the two are anti-correlated (*R* = - 0.48, *p* = - 4.35 10^-59^). In other words, structure and function closely correspond in unimodal cortex, but diverge as one moves up the hierarchy. At the apex of the hierarchy (transmodal cortex), there is much less correspondence between structural and functional connection profiles.

### Alternative predictors and individual participants

As a final step, we ask two important questions. First, how sensitive are the overall results to choice of predictors? Thus far, we focused on two canonical network metrics, one related to shortest path routing and the other related to diffusion. The relative contribution of each variable, estimated using stepwise regression, is shown in Fig. S3. Though theoretically driven, the choice of these two measures is arbitrary and there exist several other network-theoretic statistics that also capture the potential for two nodes to exchange signals with each other [3], including alternative forms of diffusion [26, 51, 53], contagion [49], parallel exchange via path ensembles [2] and navigation [67].

We therefore repeated the analysis shown above using a multilinear model with a greater number of predictors (Euclidean distance, path length and communicability as before, and adding search information, path transitivity [26, 62, 69, 73]). We note two key results. First, the overall model fit does not change appreciably, with the mean *R*^2^ = 0.33 (median *R*^2^ = 0.33, and standard deviation = 0.12). This is unsurprising, given the well-known multicolinearity among graph measures [55]. More importantly, the spatial distribution of *R*^2^ values is highly correlated those produced by a multilinear model with fewer predictors (*R* = 0.98, *p <* 10^−5^), suggesting little practical benefit for including additional predictors.

The second question is to what extent can comparable effects be observed in individual participants? In an effort to amplify the signal to noise ratio we initially performed all analyses on group-representative structural and functional networks, and it is unclear whether the systematic divergence between structure and function is robust across individuals. We therefore fit a multilinear model to each individual participant and estimated regional *R*^2^ values as before. We then correlated the individual-level *R*^2^ pattern with the group-level *R*^2^ pattern. The individual-to-group correlation *R* is moderate (mean *R* = 0.33, median *R* = 0.32, 95% CI [0.07 0.50]). but statistically significant (*p <* 0.05 in 39/40 participants), suggesting considerable consistency across individuals.

## DISCUSSION

The present report demonstrates variation in the extent to which structure and function correspond in human cortical networks. The relationship between structural and functional connection profiles appears to follow an overarching cognitive-representational and cytoarchitectural hierarchy, becoming increasingly untethered as one moves towards transmodal cortex at the apex.

### Localized structure-function relationships

Our results contribute to a growing effort to understand structure and function from a more localized perspective. There is a rich literature on predicting function from structure at the whole-network level, including direct edge-to-edge comparisons [30, 68], multivariate statistical models [44, 47], network-theoretic models [1, 14, 26, 49, 67] and biophysical models [8, 19, 65]. We find that the relationship may not be uniform throughout the whole network, but may instead vary across brain regions. This is consistent with the notion that individual areas possess distinct connectional [57, 71] and spectral activation “fingerprints” [38]. High-density precision mapping studies suggest that functional organization and regional boundaries may also be highly individualized [27, 40].

For the present analysis we chose two predictors that cover the extremes of a putative communication spectrum [3], one reflecting routing of information and one reflecting diffusion. The extent to which signaling is centralized or decentralized is an exciting open question [28, 67]. For instance, individual areas may broadcast information differently from one another, while large-scale systems may utilize different forwarding protocols or frequency channels [23]. Our results open the possibility that communication mechanisms may be multiplexed, with multiple protocols operating in parallel [26].

It is noteworthy that simple Euclidean distance was a powerful predictor. The probability of structural connectivity [32, 60], and the magnitude of functional connectivity between areas both decrease with spatial separation [50, 64]. Indeed, many topological attributes of brain networks can be accounted for by simple generative mechanisms that minimize interareal wiring cost [61, 70] (but see also [4, 79]). Our results are consistent with this notion, showing that the spatial embedding of brain regions is the most informative predictor of their functional interactions.

### Functional and cytoarchitectonic hierarchies

More generally, our findings contribute to an emerging literature that emphasizes macroscopic spatial gradients as a primary organizing principle [34, 36, 39, 46]. Smooth variation across cortex has been observed in gene expression [11], cytoarchitecture [24], myeloarchitecture [33], cortical thickness [81], structural connectivity [9] and functional connectivity [42]. The increasing complexity of cortical microcircuitry along this hierarchical gradient, ranging from primary sensory to transmodal cortex, is thought to engender increasingly integrative internal representations and functions.

Our findings suggest that a consequence of hierarchical microscale organization is a gradual decoupling of macroscale structure and function. In primary sensory areas we find a close correspondence between structural and functional connection profiles, but at the apex of the hierarchy - corresponding to the default mode and salience networks - the two diverge considerably. The polyfunctional hubs that occupy this end of the gradient are thus more likely to participate in multiple networks and explore a wider dynamic repertoire [78, 86]. How the correspondence between node-level structure and function relates to individual differences in behaviour is an exciting question for future work [43].

Why do structural and functional networks come untethered? One explanation could be that static network-theoretic metrics do not adequately capture the dynamic mechanisms that give rise to functional interactions. For instance, we have previously suggested that the network embedding of polysensory association areas places them in an optimal position to simultaneously receive signals originating from multiple sources across the network [49, 83]. Thus, extensive mixing of diverse signals at top of the hierarchy may engender less predictable functional relationships and wider discrepancy between structure and function.

An alternative explanation is that the increasing complexity of local microcircuitry contributes to the overall signal variance in transmodal cortex. In particular, the shifts in structure-function relationships mirror patterns of laminar differentiation [46, 56]. In primary areas with strong differentiation there is a strong correspondence between structure function, while in transmodal cortex - with weaker laminar differentiation - the structure-function relationship is also weaker. In a recent modeling study, Wang and colleagues allowed microscale-related parameters of a biophysical model to differ between brain regions [82]. The best-fitting model was characterized by strong recurrent connections and excitatatory subcortical input in sensorimotor regions; conversely, default network regions had weak recurrent connections and excitatory subcortical inputs [82]. Complementary results were reported by Demirta’s and colleagues, who found that biophysical models could be fitted to functional connectivity much better if they were informed by hierarchical heterogeneity, estimated from T1w/T2w ratios [20]. Thus, a richer local cytoarchitecture in transmodal cortex - supporting increasingly autonomous and spontaneous dynamics - may potentially render macroscale structural metrics overall less effective in predicting functional interactions [16].

### Methodological considerations

The present results are subject to several important methodological limitations and considerations. First, structural connectivity is estimated using streamline tractography on diffusion weighted imaging, a method known to be susceptible to systematic false positives and false negatives [17, 41, 85]. In addition, previous reports have found evidence that functional connectivity may also be more variable in heteromodal or transmodal cortex [54]. Although the current results are derived using high-resolution diffusion spectrum imaging and in a group-consensus networks, it is nevertheless possible that there is systematic underrepresentation or mischaracterization of structural or functional connectivity in transmodal cortex, manifesting as a lower correspondence between structural and functional connectivity.

A second concern is that our results are based on parcellated data, a methodological approach that assumes that brain regions can be mapped to identical spatial locations in every participant. Recent evidence from precision mapping studies, using repeated measurements in single individuals, suggests that functional boundaries can systematically vary across individuals and that this is particularly true in higher-order, transmodal cortex [27, 40].

Finally, it is important to acknowledge that the present multilinear model(s) violate a basic assumption of regression models, namely that the observations (regional connection profiles) are not independent. Each observation represents a dyadic (*i, j*) relationship that is drawn from a graph that represents the brain, a system we know to be spatially contiguous and assume to be connected. The expected effect is that parameter estimates and goodness-of-fit metrics will therefore be biased. For this reason, we only use structure-function *R*^2^ as a relative metric to compare the correspondence of structure and function across a set of nodes, each of which is estimated under the same conditions.

## METHODS

### Data acquisition

We performed all analyses in two datasets. The main (discovery) dataset was collected at the Department of Radiology, University Hospital Center and University of Lausanne, (LAU; N=40). We also included a replication cohort from the Human Connectome Project (HCP; N=215) [76]. Structural connectivity was reconstructed from diffusion-weighted imaging: diffusion spectrum imaging (DSI) for LAU, high angular resolution diffusion imaging (HARDI) for HCP. Although dataset LAU had fewer participants, we selected it as the discovery dataset because of the quality of the DSI sequence. Below we describe the acquisition, processing and connectome reconstruction procedure for each dataset in more detail.

#### LAU

A total of N = 40 healthy young adults (16 females, 25.3 ± 4.9 years old) were scanned at the Department of Radiology, University Hospital Center and University of Lausanne. The scans were performed in 3-Tesla MRI scanner (Trio, Siemens Medical, Germany) using a 32-channel head-coil. The protocol included (1) a magnetization-prepared rapid acquisition gradient echo (MPRAGE) sequence sensitive to white/gray matter contrast (1 mm in-plane resolution, 1.2 mm slice thickness), (2) a diffusion spectrum imaging (DSI) sequence (128 diffusion-weighted volumes and a single b0 volume, maximum b-value 8000 s*/*mm^2^, 2.2 × 2.2 × 3.0 mm voxel size), and (3) a gradient echo EPI sequence sensitive to BOLD contrast (3.3 mm in-plane resolution and slice thickness with a 0.3 mm gap, TR 1920 ms, resulting in 280 images per participant). Participants were not subject to any overt task demands during the fMRI scan.

#### HCP

A total of N = 215 healthy young adults (112 females, 29.7 ± 3.4 years old) were scanned as part of the HCP Q3 release [25, 76]. MRI data were acquired on the HCP’s custom 3-Tesla Siemens Skyra with a 32-channel head coil. The protocol included (1) a 3D-MPRAGE sequence, (2) a high angular resolution diffusion imaging (HARDI) sequence, and (3) a multi-band accelerated 2D-BOLD EPI sequence sensitive to BOLD contrast. For more details regarding the acquisition protocol see [25, 76].

### Structural network reconstruction

Grey matter was parcellated into 68 cortical nodes according to the Desikan-Killiany atlas [21]. These regions of interest were then further divided into four additional, increasingly finer-grained resolutions, comprising 114, 219, 448 and 1000 approximately equally-sized nodes [12]. Structural connectivity was estimated for individual participants using deterministic streamline tractography. The procedure was implemented in the Connectome Mapping Toolkit [15], initiating 32 streamline propagations per diffusion direction for each white matter voxel.

To mitigate concerns about inconsistencies in reconstruction of individual participant connectomes [35, 72], as well as the sensitive dependence of network measures on false positives and false negatives [85], we adopted a group-consensus approach [6, 17, 61]. In constructing a consensus adjacency matrix, we sought to preserve (a) the density and (b) the edge length distribution of the individual participants matrices [4, 6, 49].

We first collated the extant edges in the individual participant matrices and binned them according to length. The number of bins was determined heuristically, as the square root of the mean binary density across participants. The most frequently occurring edges were then selected for each bin. If the mean number of edges across participants in a particular bin is equal to *k*, we selected the *k* edges of that length that occur most frequently across participants. To ensure that inter-hemispheric edges are not under-represented, we carried out this procedure separately for inter- and intra-hemispheric edges. The binary densities for the final whole-brain matrices were 28.1%, 20.3%, 12.0%, 5.9% and 2.4% for resolutions 1 to 5, respectively.

### Functional network reconstruction

Functional MRI data were pre-processed using procedures designed to facilitate subsequent network exploration [59]. FMRI volumes were corrected for physiological variables, including regression of white matter, cerebrospinal fluid, as well as motion (three translations and three rotations, estimated by rigid body co-registration). BOLD time series were then subjected to a lowpass filter (temporal Gaussian filter with full width half maximum equal to 1.92 s). The first four time points were excluded from subsequent analysis to allow the time series to stabilize. Motion scrubbing was performed as described by Power and colleagues [59]. The data were parcellated according to the same atlas used for structural networks [12].

A group-average functional connectivity matrix was constructed from the fMRI BOLD time series by concatenating the regional time series from all participants and estimating a single correlation matrix. To threshold this matrix, we sampled at random 276 points from the concatenated times series and re-calculated a full correlation matrix from these points (1,000 repetitions). From these bootstrapped samples, we estimated confidence intervals for the correlation magnitude between every pair of brain regions. Pairs whose correlation was consistently positive or negative across the 1000 samples were retained (along with the sign and weight of the correlation) as putative functional connections.

### Multilinear model

A multiple regression model was used to predict the functional connection profile of every node using a set of geometric and structural connection profile predictors of the same node (Fig. 1). The predictors were (1) the Euclidean distance between node centroids, (2) path length between nodes and (3) communicability between nodes. Path length and communicability were both estimated from the binarized structural connectome. Path length refers to the shortest contiguous sequence of edges between two nodes. Communicability (*C_ij_*) between two nodes *i* and *j* is defined as the weighted sum of all paths and walks between those nodes [22]. For a binary adjacency matrix **A**, communicability is defined as

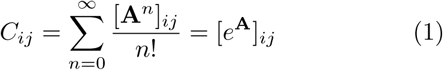

with walks of length *n* normalized by *n*!, ensuring that shorter, more direct walks contribute more than longer walks. Both metrics were implemented in the Brain Connectivity Toolbox (https://sites.google.com/site/bctnet/) [63].

The regression model was then constructed for each node *i*

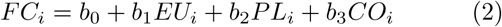

where the response variable *F C_i_* is the set of functional connections between *i* and all other nodes, and the predictor variables are the Euclidean distance (*EU_i_*), structural path length (*P L_i_*) and structural communicability (*CO_i_*) between *i* and all other nodes in the network. The regression coefficients *b*_1_, *b*_2_ and *b*_3_, as well as the intercept *b*_0_ were then solved by ordinary least squares (function *fitlm.m* in MATLAB 2016a).

### Diffusion map embedding

Diffusion map embedding is a nonlinear dimensionality reduction algorithm [13]. The algorithm seeks to project a set of embeddings into a lower-dimensional Euclidean space. Briefly, the similarity matrix among a set of points (in our case, the correlation matrix representing functional connectivity) is treated as a graph, and the goal of the procedure is to identify points that are proximal to one another on the graph. In other words, two points are close together if there are many relatively short paths connecting them. A diffusion operator, representing an ergodic Markov chain on the network, is formed by taking the normalized graph Laplacian of the matrix. The new coordinate space is described by the eigenvectors of the diffusion operator. In keeping with previous reports that applied the method to functional networks, we set the diffusion rate α*α*= 0.5 [18, 42], which approximates the Fokker-Planck diffusion. The procedure was implemented using the Dimensionality Reduction Tool-box (https://lvdmaaten.github.io/drtoolbox/) [75].

## Acknowledgments

The authors thank Dr. Alessandra Griffa for collecting, preprocessing and sharing the Lausanne dataset. The authors also thank Dr. František Vaša for contributing the parcellated von Economo cytoarchitectural atlas originally developed by Scholtens and colleagues [66]. This research was undertaken thanks in part to funding from the Canada First Research Excellence Fund, awarded to McGill University for the Healthy Brains for Healthy Lives initiative. BM acknowledges support from the Natural Sciences and Engineering Research Council of Canada (NSERC Discovery Grant RGPIN #017-04265), from the Canada Research Chairs Program and from the Fonds de recherche du Québec - Santé (Chercheur Boursier).

---

**Figure S1.**
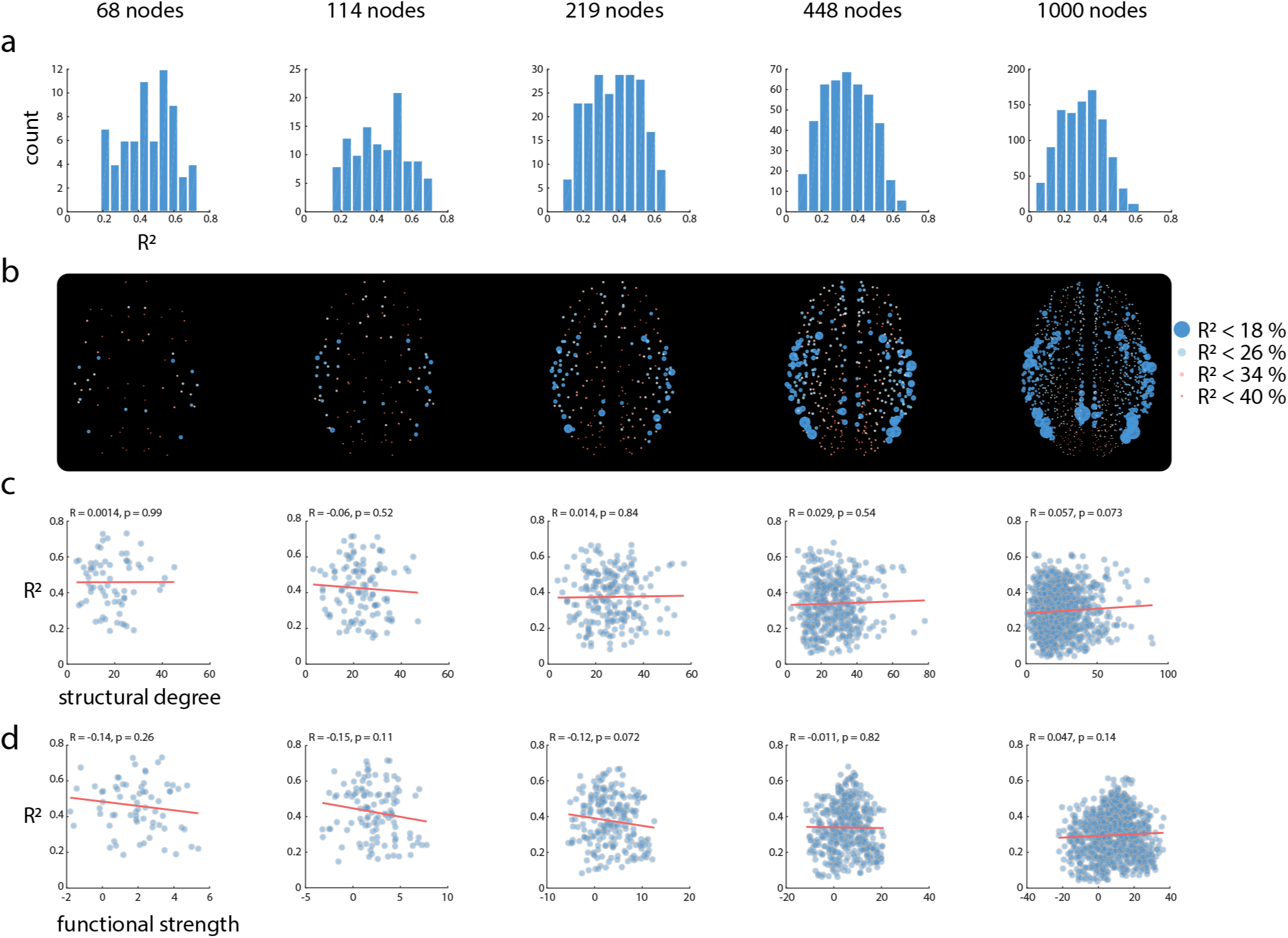
Stability of main results across five parcellations. The results shown in Fig. 2 are repeated for five anatomical parcellations provided by Cammoun and colleagues [12], featuring 68, 114, 219, 448 and 1000 cortical nodes. (a) Histograms of node-wise *R*^2^ values from the structure-function multilinear model. (b) Spatial distributions of *R*^2^ values. (c) Correlations between node structural degree and *R*^2^ values. (d) Correlations between node functional strength and *R*^2^ values.

**Figure S2.**
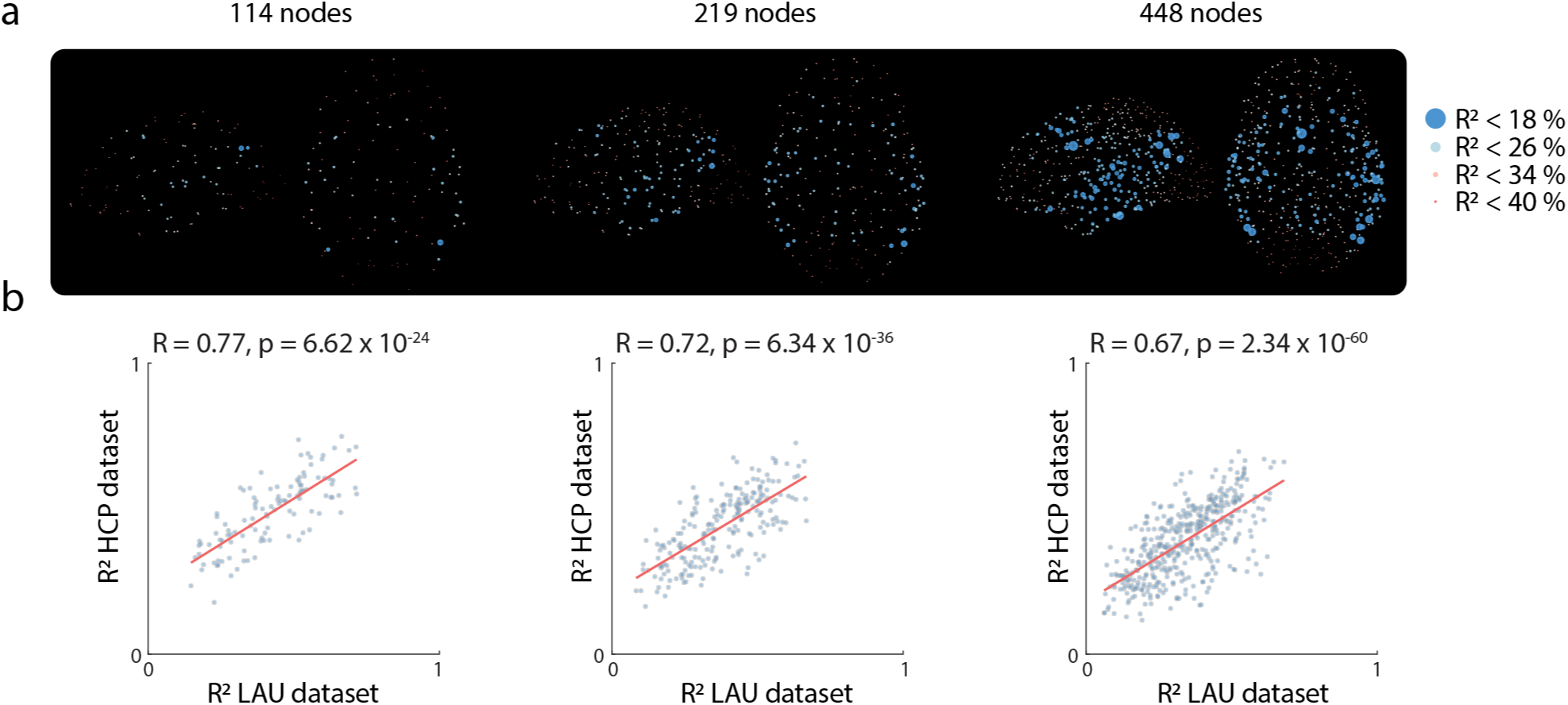
Replication dataset. To determine whether the results are replicable, we fit the multilinear model in an independently-collected dataset. (a) The spatial distribution of *R*^2^ at 3 different parcellation resolutions. Nodes with smaller structure-function *R*^2^ values are indicated by larger circles and colder colours. (b) The correlation between node-wise *R*^2^ in the discovery dataset (Lausanne dataset; LAU; *N* = 40) and the validation dataset (Human Connectome Project dataset; HCP; *N* = 215).

**Figure S3.**
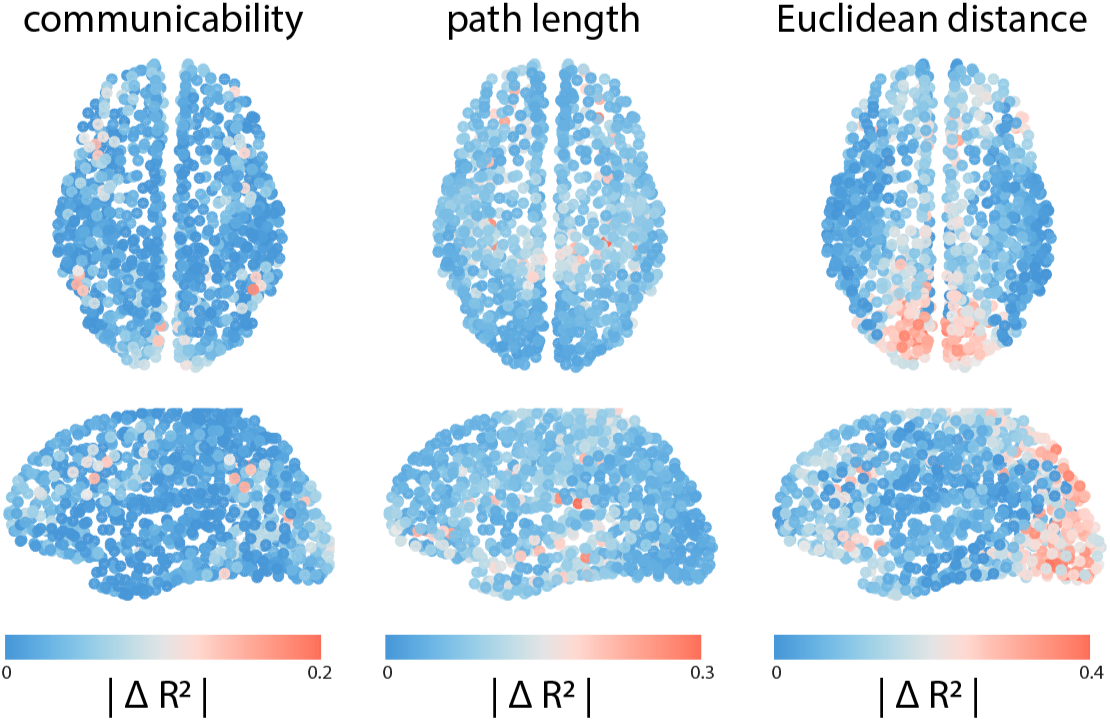
Variable importance. To assess the contribution of individual independent variables in the multilinear model, we first estimate the structure-function *R*^2^ with all variables, as well as the *R*^2^ when individual variables are removed. The contribution of a variable is quantified as the decrease in model fit (∆*R*^2^) following removal. Node-wise ∆*R*^2^ is displayed for communicability, path length and Euclidean distance. All values were negative, so absolute values are shown for simplicity.

## References

[1] Abdelnour, F., Voss, H. U., and Raj, A. (2014). Network diffusion accurately models the relationship between structural and functional brain connectivity networks. NeuroImage, 90:335–347.

[2] Avena-Koenigsberger, A., Mišić, B., Hawkins, R. X., Griffa, A., Hagmann, P., Goñi, J., and Sporns, O. (2017). Path ensembles and a tradeoff between communication efficiency and resilience in the human connectome. Brain Struct and Funct, 222(1):603–618.

[3] Avena-Koenigsberger, A., Misic, B., and Sporns, O. (2018). Communication dynamics in complex brain networks. Nat Rev Neurosci, 19(1):17.

[4] Betzel, R. F., Avena-Koenigsberger, A., Goñi, J., He, Y., De Reus, M. A., Griffa, A., Vértes, P. E., Mišic, B., Thiran, J.-P., Hagmann, P., et al. (2016). Generative models of the human connectome. NeuroImage, 124:1054–1064.

[5] Betzel, R. F. and Bassett, D. S. (2018). Specificity and robustness of long-distance connections in weighted, inter-areal connectomes. Proc Natl Acad Sci USA, 115:E4880–E4889.

[6] Betzel, R. F., Griffa, A., Hagmann, P., and Mišić, B. (2018). Distance-dependent consensus thresholds for generating group-representative structural brain networks. Network Neuroscience, pages 1–22.

[7] Betzel, R. F., Medaglia, J. D., Kahn, A. E., Soffer, J., Schonhaut, D. R., and Bassett, D. S. (2017). Inter-regional ecog correlations predicted by communication dynamics, geometry, and correlated gene expression. arXiv preprint arXiv:1706.06088.

[8] Breakspear, M. (2017). Dynamic models of large-scale brain activity. Nat Neurosci, 20(3):340.

[9] Buckner, R. L. and Margulies, D. S. (2018). Macroscale cortical organization and a default-like transmodal apex network in the marmoset monkey. bioRxiv, page 415141.

[10] Bullmore, E. and Sporns, O. (2009). Complex brain networks: graph theoretical analysis of structural and functional systems. Nat Rev Neurosci, 10(3):186.

[11] Burt, J. B., Demirta’s, M., Eckner, W. J., Navejar, N. M., Ji, J. L., Martin, W. J., Bernacchia, A., Anticevic, A., and Murray, J. D. (2018). Hierarchy of transcriptomic specialization across human cortex captured by structural neuroimaging topography. Nat Neurosci, 21(9):1251.

[12] Cammoun, L., Gigandet, X., Meskaldji, D., Thiran, J. P., Sporns, O., Do, K. Q., Maeder, P., Meuli, R., and Hagmann, P. (2012). Mapping the human connectome at multiple scales with diffusion spectrum mri. J Neurosci Meth, 203(2):386–397.

[13] Coifman, R. R., Lafon, S., Lee, A. B., Maggioni, M., Nadler, B., Warner, F., and Zucker, S. W. (2005). Geometric diffusions as a tool for harmonic analysis and structure definition of data: Diffusion maps. Proc Natl Acad Sci USA, 102(21):7426–7431.

[14] Crofts, J. J. and Higham, D. J. (2009). A weighted communicability measure applied to complex brain networks. J Roy Soc Interface, pages rsif–2008.

[15] Daducci, A., Gerhard, S., Griffa, A., Lemkaddem, A., Cammoun, L., Gigandet, X., Meuli, R., Hagmann, P., and Thiran, J.-P. (2012). The connectome mapper: an open-source processing pipeline to map connectomes with mri. PLoS ONE, 7(12):e48121.

[16] Dagher, A., Clark, C. A., Shafiei, G., Zeighami, Y., Mii, B., Nagano-Saito, A., Leyton, M., and Coull, J. T. (2018). Dopamine Signaling Modulates the Stability and Integration of Intrinsic Brain Networks. Cereb Cortex, 29(1):397–409.

[17] de Reus, M. A. and van den Heuvel, M. P. (2013). Estimating false positives and negatives in brain networks. NeuroImage, 70:402–409.

[18] de Wael, R. V., Lariviére, S., Caldairou, B., Hong, S.-J., Margulies, D. S., Jefferies, E., Bernasconi, A., Smallwood, J., Bernasconi, N., and Bernhardt, B. C. (2018). Anatomical and microstructural determinants of hippocampal subfield functional connectome embedding. Proc Natl Acad Sci USA, 115(40):10154–10159.

[19] Deco, G., Jirsa, V., McIntosh, A. R., Sporns, O., and Kötter, R. (2009). Key role of coupling, delay, and noise in resting brain fluctuations. Proc Natl Acad Sci USA, pages pnas–0901831106.

[20] Demirtas, M., Burt, J. B., Helmer, M., Ji, J. L., Adkinson, B. D., Glasser, M. F., Van Essen, D. C., Sotiropoulos, S. N., Anticevic, A., and Murray, J. D. (2010). Hierarchical heterogeneity across human cortex shapes large-scale neural dynamics. Neuron, page 341966.

[21] Desikan, R. S., Ségonne, F., Fischl, B., Quinn, B. T., Dickerson, B. C., Blacker, D., Buckner, R. L., Dale, M., Maguire, R. P., Hyman, B. T., et al. (2006). An automated labeling system for subdividing the human cerebral cortex on mri scans into gyral based regions of interest. NeuroImage, 31(3):968–980.

[22] Estrada, E. and Hatano, N. (2008). Communicability in complex networks. Phys Rev E, 77(3):036111.

[23] Fries, P. (2005). A mechanism for cognitive dynamics: neuronal communication through neuronal coherence. Trends Cogn Sci, 9(10):474–480.

[24] Fulcher, B. D., Murray, J. D., Zerbi, V., and Wang, X.-J. (2019). Multimodal gradients across mouse cortex. Proc Natl Acad Sci USA, pages –.

[25] Glasser, M. F., Sotiropoulos, S. N., Wilson, J. A., Coalson, T. S., Fischl, B., Andersson, J. L., Xu, J., Jbabdi, S., Webster, M., Polimeni, J. R., et al. (2013). The minimal preprocessing pipelines for the human connectome project. NeuroImage, 80:105–124.

[26] Goñi, J., van den Heuvel, M. P., Avena-Koenigsberger, A., de Mendizabal, N. V., Betzel, R. F., Griffa, A., Hagmann, P., Corominas-Murtra, B., Thiran, J.-P., and Sporns, O. (2014). Resting-brain functional connectivity predicted by analytic measures of network communication. Proc Natl Acad Sci USA, 111(2):833–838.

[27] Gordon, E. M., Laumann, T. O., Gilmore, A. W., Newbold, D. J., Greene, D. J., Berg, J. J., Ortega, M., Hoyt-Drazen, C., Gratton, C., Sun, H., et al. (2017). Precision functional mapping of individual human brains. Neuron, 95(4):791–807.

[28] Graham, D. and Rockmore, D. (2011). The packet switching brain. J Cogn Neurosci, 23(2):267–276.

[29] Hilgetag, C. C. and Kaiser, M. (2004). Clustered organization of cortical connectivity. Neuroinformatics, 2(3):353–360.

[30] Honey, C., Sporns, O., Cammoun, L., Gigandet, X., Thiran, J.-P., Meuli, R., and Hagmann, P. (2009). Predicting human resting-state functional connectivity from structural connectivity. Proc Natl Acad Sci USA, 106(6):2035–2040.

[31] Honey, C. J., Kötter, R., Breakspear, M., and Sporns, O. (2007). Network structure of cerebral cortex shapes functional connectivity on multiple time scales. Proc Natl Acad Sci USA, 104(24):10240–10245.

[32] Horvát, S., Gămănu, R., Ercsey-Ravasz, M., Magrou, L., Gămănu, B., Van Essen, D. C., Burkhalter, A., Knoblauch, K., Toroczkai, Z., and Kennedy, H. (2016). Spatial embedding and wiring cost constrain the functional layout of the cortical network of rodents and primates. PLoS biology, 14(7):e1002512.

[33] Huntenburg, J. M., Bazin, P.-L., Goulas, A., Tardif, C. L., Villringer, A., and Margulies, D. S. (2017). A systematic relationship between functional connectivity and intracortical myelin in the human cerebral cortex. Cereb Cortex, 27(2):981–997.

[34] Huntenburg, J. M., Bazin, P.-L., and Margulies, D. S. (2018). Large-scale gradients in human cortical organization. Trends Cogn Sci, 22(1):21–31.

[35] Jones, D., Knösche, T., and Turner, R. (2013). White matter integrity, fiber count, and other fallacies: the do’s and don’ts of diffusion mri. NeuroImage, 73:239–254.

[36] Jones, E. and Powell, T. (1970). An anatomical study of converging sensory pathways within the cerebral cortex of the monkey. Brain, 93(4):793–820.

[37] Kaiser, M. and Hilgetag, C. C. (2006). Nonoptimal component placement, but short processing paths, due to long-distance projections in neural systems. PLoS Comput Biol, 2(7):e95.

[38] Keitel, A. and Gross, J. (2016). Individual human brain areas can be identified from their characteristic spectral activation fingerprints. PLoS Biol, 14(6):e1002498.

[39] Larivière, S., Vos de Wael, R., Paquola, C., Hong, S.-J., Mišić, B., Bernasconi, N., Bernasconi, A., Bonilha, L., and Bernhardt, B. C. (2018). Microstructure-informed connectomics: enriching large-scale descriptions of healthy and diseased brains. Brain Conn.

[40] Laumann, T. O., Gordon, E. M., Adeyemo, B., Snyder, A. Z., Joo, S. J., Chen, M.-Y., Gilmore, A. W., McDermott, K. B., Nelson, S. M., Dosenbach, N. U., et al. (2015). Functional system and areal organization of a highly sampled individual human brain. Neuron, 87(3):657–670.

[41] Maier-Hein, K. H., Neher, P. F., Houde, J.-C., Côté, M.-A., Garyfallidis, E., Zhong, J., Chamberland, M., Yeh, F.-C., Lin, Y.-C., Ji, Q., et al. (2017). The challenge of mapping the human connectome based on diffusion tractography. Nat Commun, 8(1):1349.

[42] Margulies, D. S., Ghosh, S. S., Goulas, A., Falkiewicz, M., Huntenburg, J. M., Langs, G., Bezgin, G., Eickhoff, S. B., Castellanos, F. X., Petrides, M., Jeffries, E., and Smallwood, J. (2016). Situating the default-mode network along a principal gradient of macroscale cortical organization. Proc Natl Acad Sci USA, 113(44):12574–12579.

[43] Medaglia, J. D., Huang, W., Karuza, E. A., Kelkar, A., Thompson-Schill, S. L., Ribeiro, A., and Bassett, D. S. (2018). Functional alignment with anatomical networks is associated with cognitive flexibility. Nat Hum Behav, 2(2):156.

[44] Messé, A., Rudrauf, D., Benali, H., and Marrelec, G. (2014). Relating structure and function in the human brain: relative contributions of anatomy, stationary dynamics, and non-stationarities. PLoS Comput Biol, 10(3):e1003530.

[45] Messé, A., Rudrauf, D., Giron, A., and Marrelec, G. (2015). Predicting functional connectivity from structural connectivity via computational models using mri: an extensive comparison study. NeuroImage, 111:65–75.

[46] Mesulam, M.-M. (1998). From sensation to cognition. Brain: a journal of neurology, 121(6):1013–1052.

[47] Mišić, B., Betzel, R. F., De Reus, M. A., Van Den Heuvel, M. P., Berman, M. G., McIntosh, A. R., and Sporns, O. (2016). Network-level structure-function relationships in human neocortex. Cereb Cortex, 26(7):3285–3296.

[48] Mišić, B., Betzel, R. F., Griffa, A., de Reus, M. A., He, Y., Zuo, X.-N., van den Heuvel, M. P., Hagmann, P., Sporns, O., and Zatorre, R. J. (2018). Network-based asymmetry of the human auditory system. Cereb Cortex, 28(7):2655–2664.

[49] Mišić, B., Betzel, R. F., Nematzadeh, A., Goñi, J., Griffa, A., Hagmann, P., Flammini, A., Ahn, Y.-Y., and Sporns, O. (2015). Cooperative and competitive spreading dynamics on the human connectome. Neuron, 86(6):1518–1529.

[50] Mišić, B., Fatima, Z., Askren, M. K., Buschkuehl, M., Churchill, N., Cimprich, B., Deldin, P. J., Jaeggi, S., Jung, M., Korostil, M., et al. (2014a). The functional connectivity landscape of the human brain. PLoS One, 9(10):e111007.

[51] Mišić, B., Goñi, J., Betzel, R. F., Sporns, O., and McIntosh, A. R. (2014b). A network convergence zone in the hippocampus. PLoS Comput Biol, 10(12):e1003982.

[52] Mišić, B. and Sporns, O. (2016). From regions to connections and networks: new bridges between brain and behavior. Curr Opin Neurobiol, 40:1–7.

[53] Mišić, B., Sporns, O., and McIntosh, A. R. (2014c). Communication efficiency and congestion of signal traffic in large-scale brain networks. PLoS Comput Biol, 10(1):e1003427.

[54] Mueller, S., Wang, D., Fox, M. D., Yeo, B. T., Sepulcre, J., Sabuncu, M. R., Shafee, R., Lu, J., and Liu, H. (2013). Individual variability in functional connectivity architecture of the human brain. Neuron, 77(3):586–595.

[55] Oldham, S., Fulcher, B., Parkes, L., Arnatkeviciute, A., Suo, C., and Fornito, A. (2018). Consistency and differences between centrality metrics across distinct classes of networks. arXiv preprint arXiv:1805.02375.

[56] Paquola, C., de Wael, R. V., Wagstyl, K., Bethlehem, R., Seidlitz, J., Hong, S.-J., Bullmore, E., Evans, A., Misic, B., Margulies, D., et al. (2018). Dissociations between microstructural and functional hierarchies within regions of transmodal cortex. bioRxiv, page 488700.

[57] Passingham, R. E., Stephan, K. E., and Kötter, R. (2002). The anatomical basis of functional localization in the cortex. Nat Rev Neurosci, 3(8):606.

[58] Paus, T., Pesaresi, M., and French, L. (2014). White matter as a transport system. Neuroscience, 276:117–125.

[59] Power, J. D., Barnes, K. A., Snyder, A. Z., Schlaggar, B. L., and Petersen, S. E. (2012). Spurious but systematic correlations in functional connectivity mri networks arise from subject motion. NeuroImage, 59(3):2142–2154.

[60] Roberts, J. A., Perry, A., Lord, A. R., Roberts, G., Mitchell, P. B., Smith, R. E., Calamante, F., and Breakspear, M. (2016). The contribution of geometry to the human connectome. NeuroImage, 124:379–393.

[61] Roberts, J. A., Perry, A., Roberts, G., Mitchell, P. B., and Breakspear, M. (2017). Consistency-based thresholding of the human connectome. NeuroImage, 145:118–129.

[62] Rosvall, M., Grönlund, A., Minnhagen, P., and Sneppen, K. (2005). Searchability of networks. Physical Review E, 72(4):046117.

[63] Rubinov, M. and Sporns, O. (2010). Complex network measures of brain connectivity: uses and interpretations. NeuroImage, 52(3):1059–1069.

[64] Salvador, R., Suckling, J., Coleman, M. R., Pickard, J. D., Menon, D., and Bullmore, E. (2005). Neurophysiological architecture of functional magnetic resonance images of human brain. Cereb Cortex, 15(9):1332–1342.

[65] Sanz-Leon, P., Knock, S. A., Spiegler, A., and Jirsa, V. K. (2015). Mathematical framework for large-scale brain network modeling in the virtual brain. NeuroImage, 111:385–430.

[66] Scholtens, L. H., de Reus, M. A., de Lange, S. C., Schmidt, R., and van den Heuvel, M. P. (2018). An mri von economo–koskinas atlas. NeuroImage, 170:249–256.

[67] Seguin, C., van den Heuvel, M. P., and Zalesky, A. (2018). Navigation of brain networks. Proc Natl Acad Sci USA, 115:6297–6302.

[68] Shen, K., Mišić, B., Cipollini, B. N., Bezgin, G., Buschkuehl, M., Hutchison, R. M., Jaeggi, S. M., Kross, E., Peltier, S. J., Everling, S., et al. (2015). Stable long-range interhemispheric coordination is supported by direct anatomical projections. Proc Natl Acad Sci USA, page 201503436.

[69] Sneppen, K., Trusina, A., and Rosvall, M. (2005). Hide-and-seek on complex networks. Europhys Lett, 69(5):853.

[70] Song, H. F., Kennedy, H., and Wang, X.-J. (2014). Spatial embedding of structural similarity in the cerebral cortex. Proc Natl Acad Sci USA, 111(46):16580–16585.

[71] Sporns, O. and Kötter, R. (2004). Motifs in brain networks. PLoS Biol, 2(11):e369.

[72] Thomas, C., Frank, Q. Y., Irfanoglu, M. O., Modi, P., Saleem, K. S., Leopold, D. A., and Pierpaoli, C. (2014). Anatomical accuracy of brain connections derived from diffusion mri tractography is inherently limited. Proc Natl Acad Sci USA, 111(46):16574–16579.

[73] Trusina, A., Rosvall, M., and Sneppen, K. (2005). Communication boundaries in networks. Physical review letters, 94(23):238701.

[74] van den Heuvel, M. P., Kahn, R. S., Goñi, J., and Sporns, O. (2012). High-cost, high-capacity backbone for global brain communication. Proc Natl Acad Sci USA, 109(28):11372–11377.

[75] Van Der Maaten, L., Postma, E., and Van den Herik, J. (2009). Dimensionality reduction: a comparative. J Mach Learn Res, 10:66–71.

[76] Van Essen, D. C., Smith, S. M., Barch, D. M., Behrens, T. E., Yacoub, E., Ugurbil, K., Consortium, W.-M.H., et al. (2013). The wu-minn human connectome project: an overview. NeuroImage, 80:62–79.

[77] Váša, F., Seidlitz, J., Romero-Garcia, R., Whitaker, K. J., Rosenthal, G., Vértes, P. E., Shinn, M., Alexander-Bloch, A., Fonagy, P., Dolan, R. J., et al. (2017). Adolescent tuning of association cortex in human structural brain networks. Cereb Cortex, 28(1):281–294.

[78] Vatansever, D., Menon, D. K., Manktelow, A. E., Sahakian, B. J., and Stamatakis, E. A. (2015). Default mode dynamics for global functional integration. J Neu-rosc., 35(46):15254–15262.

[79] Vértes, P. E., Alexander-Bloch, A. F., Gogtay, N., Giedd, J. N., Rapoport, J. L., and Bullmore, E. T. (2012). Simple models of human brain functional networks. Proc Natl Acad Sci USA, page 201111738.

[80] von Economo, C. F. and Koskinas, G. N. (1925). Die cytoarchitektonik der hirnrinde des erwachsenen menschen. J. Springer.

[81] Wagstyl, K., Ronan, L., Goodyer, I. M., and Fletcher, P. C. (2015). Cortical thickness gradients in structural hierarchies. NeuroImage, 111:241–250.

[82] Wang, P., Kong, R., Kong, X., Liégeois, R., Orban, C., Deco, G., van den Heuvel, M. P., and Yeo, B. T. (2019). Inversion of a large-scale circuit model reveals a cortical hierarchy in the dynamic resting human brain. Sci Adv, 5(1):eaat7854.

[83] Worrell, J., Rumschlag, J., Betzel, R., Sporns, O., and Misic, B. (2017). Optimized connectome architecture for sensory-motor integration. Net Neurosci, 1(4):415–430.

[84] Yeo, B. T., Krienen, F. M., Sepulcre, J., Sabuncu, M. R., Lashkari, D., Hollinshead, M., Roffman, J. L., Smoller, J. W., Zöllei, L., Polimeni, J. R., et al. (2011). The organization of the human cerebral cortex estimated by intrinsic functional connectivity. J Neurophysiol, 106(3):1125–1165.

[85] Zalesky, A., Fornito, A., Cocchi, L., Gollo, L. L., van den Heuvel, M. P., and Breakspear, M. (2016). Connectome sensitivity or specificity: which is more important?. NeuroImage, 142:407–420.

[86] Zhang, J., Cheng, W., Liu, Z., Zhang, K., Lei, X., Yao, Y., Becker, B., Liu, Y., Kendrick, K. M., Lu, G., et al. (2016). Neural, electrophysiological and anatomical basis of brain-network variability and its characteristic changes in mental disorders. Brain, 139(8):2307–2321.

